# Gut Phageome Analysis Reveals Disease-Specific Hallmarks in Childhood Obesity

**DOI:** 10.1101/2020.07.29.227637

**Authors:** Shirley Bikel, Gamaliel López-Leal, Fernanda Cornejo-Granados, Luigui Gallardo-Becerra, Filiberto Sánchez, Edgar Equihua-Medina, Juan Pablo Ochoa-Romo, Blanca Estela López-Contreras, Samuel Canizales-Quinteros, Adrian Ochoa Leyva

## Abstract

Changes in the composition of the human gut microbiome are recognized to have a significant association with obesity and metabolic syndrome. Mexico leads worldwide childhood-obesity rankings representing an epidemic problem for public health. To this date, it is still unclear how the gut phageome, the bacteriophage component of the virome, influences childhood obesity and obesity with metabolic syndrome. We characterized the gut phageome of 28 school-age children with healthy normal-weight (NW), obese (O), and obese with metabolic syndrome (OMS) profiles, using metagenomic sequencing of virus-like particles (VLPs) from fecal samples. Viromes derived from VLPs were mainly dominated by Caudovirales, and only Inoviridae family was significantly increased in obesity. The three groups showed a similar number of VLPs, while a significant increase in phage richness and diversity was found in obesity groups compared NW. Few phage contigs dominated the phageome composition in NW, being increased in obesity groups. Interestingly, the majority of the phageome was shared among all individuals, establishing a core and common phageome, which abundances correlated with anthropometric and biochemical traits and bacteria previously associated with obesity and metabolic syndrome. We also established a healthy core phageome shared in >80% of NW samples, with a decreased prevalence in the O and OMS groups. Our data support that changes in the gut phageome may contribute to obesity and metabolic syndrome development via bacterial dysbiosis. We consider the phageome characterization provides the basis for novel diagnostic and therapeutic strategies for managing obesity and preventing metabolic syndrome development in obese children through potential phage manipulation. To the best of our knowledge, this study represents the most in-depth sequenced dataset of human bacteriophages, demonstrating for the first time that alterations of the gut phageome characterize obesity.

## Introduction

Childhood obesity is recognized as one of the most relevant and severe health problems worldwide, it is a significant risk factor for infections such as COVID-19^1 2^, is the leading cause of adult obesity and a substantial risk for premature death^3^. It is considered a complex disease that involves genetic, environmental, and lifestyle factors, characterized by an abnormal fat accumulation due to an imbalance between energy intake and expenditure^4^. In Mexico, 17.5% of school-age children suffer obesity^5^, placing it as the country with the second-highest obesity rate in the world^6,7^.

In recent years, obesity was recognized as a state of chronic low-grade inflammation^8^, and is associated with the development of metabolic disorders including high fasting blood glucose (hyperglycemia), elevated triglycerides (hypertriglyceridemia), low levels of high-density lipoprotein (dyslipidemia), and high blood pressure (hypertension)^9^. The presence of at least three of these criteria is clinically diagnosed as metabolic syndrome^10^, a condition with high prevalence among obese children^11,12^. Notably, Mexican children are considered a high-risk ethnic group^5^ for metabolic syndrome, and its prevalence leads to an increased incidence of type 2 diabetes mellitus (T2D)^13^ and the development of cardiovascular disease (CVD)^14^ as adults. The gut microbiota is vital for human health, and alterations in its diversity and function are associated with increased energy harvest from diet, low-grade inflammation and altered adipose tissue composition^15,16^. These processes, among others, are considered as the link between gut microbiota, obesity and metabolic syndrome^17^. Our group recently demonstrated that 16S profiling and metatranscriptomic analyses showed distinct microbiota alterations among obesity and obesity with metabolic syndrome profiles^18^.

The gut virome is mainly dominated by bacteriophages (phages) and prophages^19,20^ and plays an essential role regulating microbial ecosystem and host physiology^21^ through multiple interactions and co-evolution with the co-occurring bacteriome, including horizontal gene transfer^22^, and the human immune system^23^. In this form, trans-kingdom microbiota interactions between phages, bacteria, and hosts can increase the human phenotype’s risk of disease^24–27^. Disease-specific changes of the gut virome and phageome has been reported in inflammatory bowel disease^26,28^,AIDS^29^, diabetes^30^, and malnutrition^31^. However, there are no studies addressing the role of virus in obesity and obesity with metabolic syndrome.

To address whether gut phageome plays an important role in obesity and metabolic syndrome, understanding the composition of phageome in health and disease is critical. Here, we characterized the gut phageome of healthy normal weight (NW), obese (O) and obese with metabolic syndrome (OMS) children using metagenomic sequencing of virus-like particles (VLPs) from fecal samples. Our data demonstrated for the first time that alterations of the gut phageome characterize obesity. This characterization provides the basis for novel diagnostic and therapeutic strategies for managing obesity and the prevention of metabolic syndrome in obese children. To the best of our knowledge, this is the first and most extensive study to characterize gut phageome in normal-weight and obesity and obesity with metabolic syndrome.

## Results

### The number of Virus-like particles (VLP’s) is not affected by obesity and obesity with metabolic syndrome

We used the feces collected from 28 children, ten healthy normal-weight (NW), ten obese (O), and eight obese with metabolic syndrome (OMS) children, aged 7-10 years old and paired by gender and age based on previously collected data^18^ (Extended Table 1). All individuals come from similar middle socio-economic class. Body Mass Index (BMI), Waist Circumference (WC), Triglycerides (TG), and High Density Lipoprotein (HDL) were statistically different among the NW, O, and OMS groups based on previously obtained data^18^.

The results from the transmission electron microscopy (TEM) and the fluorescence assay confirmed the correct isolation and quantification of VLPs. The majority of fluorescence corresponded to VLPs and not to bacterial sizes, suggesting that the DNA further obtained from the samples corresponded, principally, to the VLPs. Interestingly, there was no significant difference in the number of VLPs among groups, obtaining an average of 1.42×10^9^ ± 1.65 ×10^9^, 4.88×10^9^ ± 5.34×10^9^ and 3.51×10^9^ ± 4.22×10^9^ VLP’s for NW, O and OMS, respectively (Fig. 1A).

**Fig. 1.**
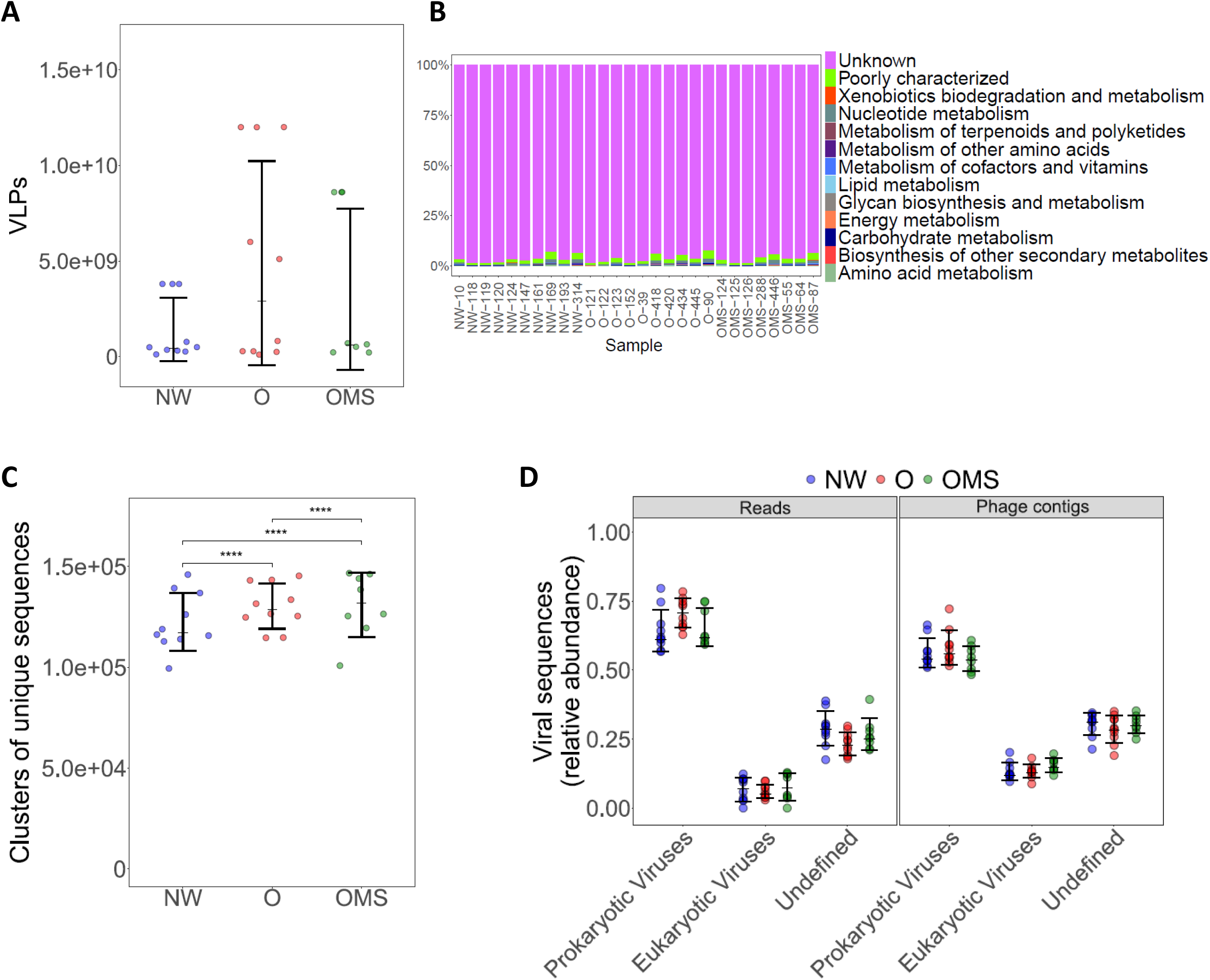
The number of VLPs and characterization of derived reads and contigs. Each panel compares the NW, O, and OMS groups. **A**. Virus-like particles (VLPs) counts. Points represent the average number of VLPs for each sample. Error bars indicate the median and interquartile range. **B**. The relative abundance of KEGG categories in VLPs derived reads. “Unknown” (purple) indicates the proportion of reads that cannot be classified. **C**. The number of clusters of unique read sequences (y-axis) per group. The points represent the average of 1000 iterations at the same sequence depth for each sample, showing each group’s distribution. Error bars represent the mean +/- SD. Statistical significance was determined by the Kruskal-Wallis test with Dunn’s correction comparing all samples to all samples. ****= p < 0.0001. **D**. The relative abundance of reads sequences and phage contigs assigned to the indicated viral taxa. Error bars represent the median and interquartile range.

### Shotgun metagenomes support viral enrichment in VLPs

Unlike previous studies, DNA extracted from VLPs was directly sequenced without prior whole-genome amplification to avoid the typical PCR amplification biases. After quality controls, we obtained an average of 4,871,075 paired-end sequences per sample (Extended Table 2). These 11.23 Gb of data, represent the most in-depth sequenced dataset of human bacteriophages to date (Extended Table 3).

To only select the sequences derived from VLPs, the reads mapped to bacterial (average of 28%), and human (average of 15%) genomes were removed for further analysis (Extended Table 2). Although, the removal of potential bacterial contamination risks also removing viral reads from a prophage state, we preferred to avoid potential bacterial DNA contamination. As a result, we obtained an average of 2,673,548 quality-filtered sequences per sample with no significant sequence depth differences among the three groups (Extended Fig. 1 and Extended Table 2). The annotation of the viral reads to the KEGG database showed that 96.2±11.89% mapped to genes with unknown function (Fig. 1B), which is in agreement with a previous virome report^32^, supporting the viral enrichment in our VLPs purification.

### Obesity and obesity with metabolic syndrome showed and increased richness of sequencing reads previous to taxonomy classification

To assess the potential viral richness before assigning a taxonomical classification, we conducted 1,000 randomly subsampled exercises of 149,000 viral reads (according to the smallest sample) and clustered them at 95% identity to generate unique clusters of sequences for each sample. These rarefactions at the same sequence depth showed a significant increase (p-value <0.0001) of unique clusters in OMS and O, followed by NW (Fig. 1C). This increased viral-reads richness in obesity groups compared to healthy normal-weight was not dependent on sequencing depth or taxonomy classification.

### Viral taxonomy of viral reads showed Caudovirales as the predominant phages in all groups

All the sequences for each group were clustered at 95% identity for each sample to remove redundant reads and to generate “unique” viral sequences. After that, we obtained a reduction of 68% ± 8%, resulting in an average of 856,825 unique sequences per sample (Extended Table 2). After mapping, only 2.95±0.95% of the unique reads matched against the viral NR Refseq protein database (Extended Fig. 2A); 66.95±6.95% of the identified sequences corresponded to prokaryotic viruses, 6.79±3.91% to eukaryotic viruses and 26.26±5.79% had undefined classification (Fig. 1D).

The taxonomy of the classified viral reads showed that the Caudovirales order represented the 94.30±0.19%, and was sub-classified as Siphoviridae (50.95±0.08%), Mioviridae (41.22±0.09%), and Podovoridae (7.82±0.03%) families. The statistical analysis showed no significant differences between these families among the three groups (Extended Fig. 2B). Considering that the variable length of the phage genomes can affect their real abundance^32^, we assembled the potential viral genomes for further in-depth sequence analysis.

### The virome assembly confirms a dominance of bacteriophages in the gut virome

We used the viral reads for *de Novo* assembly of the human gut virome. We selected contigs with a length coverage ≥80% and a depth sequencing coverage ≥1X in at least one sample to eliminate potential chimera contigs and obtained 18,602 contigs (≥500 nt; largest, 176,210 nt). In average, 58.88±13.13% of reads from each sample mapped back to these contigs, showing a relatively even contribution of all samples to the assembly. In order to decrease the probability of selecting gut eukaryotic viruses (mainly Anelloviridae family) with a reported genome size between 3-4Kb^31^ and considering the shortest genomes of gut phages are reported around ≥4Kb^33^, we eliminated contigs shorter than 4,000nt, maintaining 12,887 contigs for further analysis.

For taxonomic classification, we used the information in both the assembled contigs and their encoding proteins, obtaining 4,611 contigs as prokaryotic viruses, 1,540 as eukaryotic viruses, and 6,136 contigs with multiple hits or unknown origin. Per sample, the classification showed on average, 56.30±5.50% of prokaryotic viruses and 13.94± 2.80% of eukaryotic viruses (Fig. 1D), a similar proportion to the observed for the viral reads (Fig. 1D). Most of the prokaryotic contigs were dsDNA viruses (96%), while only 0.1% was annotated as ssDNA viruses, and the remaining 4% were unclassified bacterial viruses. We selected the 4,611 potential bacteriophages contigs (max, 176,210nt; mean, 9,347 nt) as the children’s gut phageome, hereafter mentioned as phage contigs for further analysis..

### The phageome was mainly composed of Caudovirales, and only the Inoviridae abundance was increased in obesity

As was previously observed for the sequencing reads, Caudovirales were the most abundant phage contigs in all groups (91.28±0.10%), followed by crAss-like viruses and others (Fig. 2). Within Caudovirales, the most abundant were Siphoviridae (35.28±0.02%), Myoviridae (31.33±0.03%), and Podoviridae (6.50±0.01%) families (Fig. 2). Given the importance of crAss-like phages and Microviridae in adult viromes^34,35^, we analyzed their presence and found that crAss-like phages constituted, on average, 0.64% ± 0.01% and Microviridae 0.03% ± 4.4×10^−4^ of the contigs (Extended Table 4). To compare the phage abundance among our samples, we calculated the RPKM^32^ of each phage contig, mapping back the total reads from each sample to the phage contigs. From the classified phage orders and families, the Inoviridae was the unique taxa significantly (p-value = 0.04) increased in O (RPKM_median_ = 44.35) versus NW (RPKM_median_= 4.01).

**Fig. 2:**
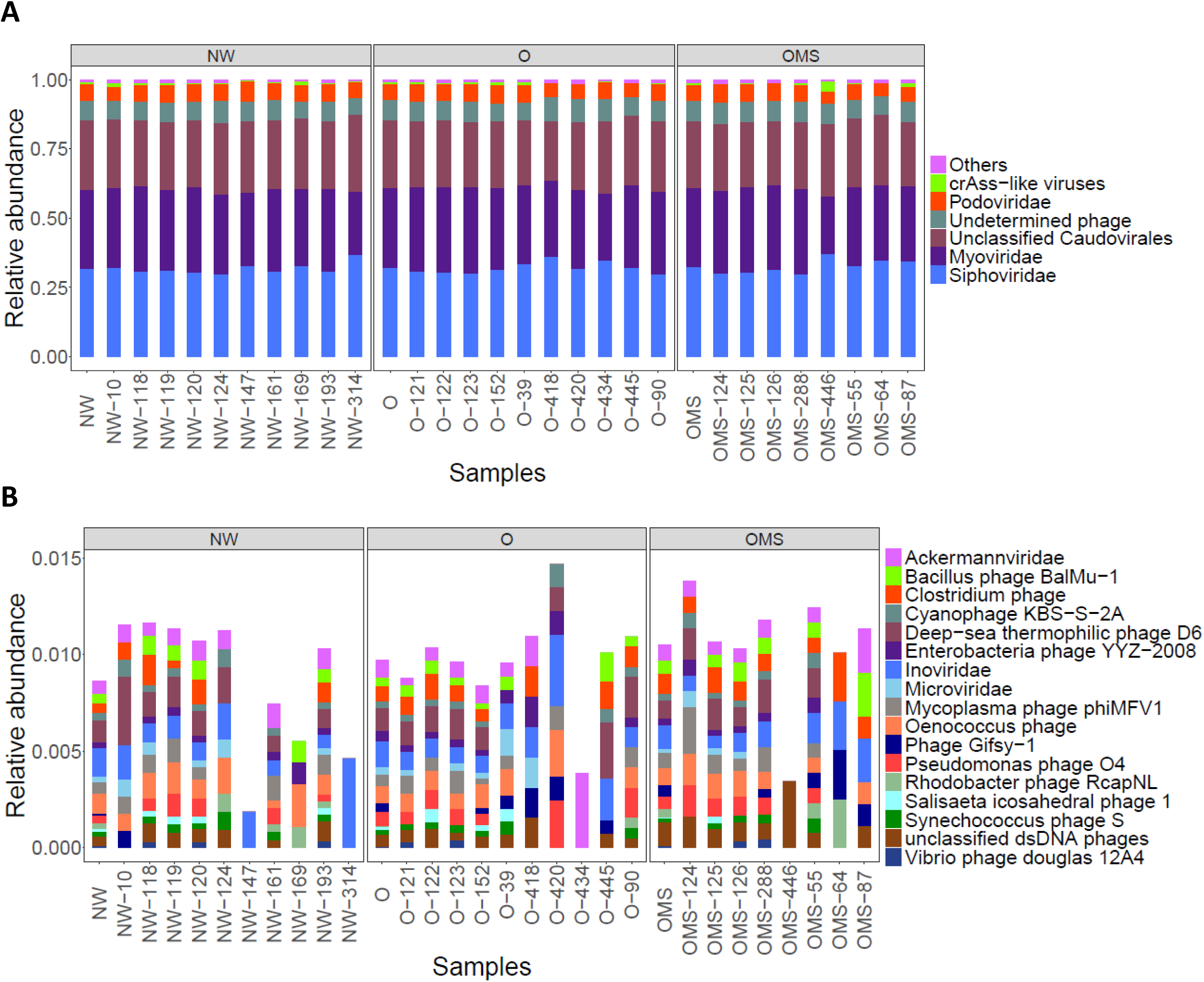
Taxonomic classification of phage contigs per group and sample. A. The relative abundance of phage contigs assigned to the indicated viral taxa. “Others” (purple) indicates the sum of the relative abundance of viral taxa showed in B. **B. The r**elative abundance of the less abundant phage contigs assigned to the indicated viral taxa.

### Over-abundant phage contigs were associated to obesity

We identified the statistically differential abundant phages using edgeR with the recruitment matrix mentioned in the previous section. This procedure is analogous to using tables of RPKM for the measurement of differentially expressed genes in RNA-seq experiments. We compared the phage contig abundances between O vs. NW, and OMS vs. NW and only selected significantly over-abundant phage contigs present in at least 30% of samples within O and OMS groups. With this cut-off, we identified phage contigs shared by the population and not individual’s specific phages. A total of 82 and 67 phage contigs were over-abundant in O and OMS, respectively, as compared to the NW, and of these, 48 (47.5%) phage contigs were shared between O and OMS (Fig. 3). The shared phage contigs belonged to Caudovirales (90%), followed by non-determined phages (6%), and Inoviridae (2%). The shared phage contigs in both two obesity groups suggest they could be associated with the core of obesity disease. It would be interesting to validate this assumption by analyzing these shared phages at functional level^36^, as well as to analyze the functional profile of the over-abundant unique phages in O and OMS groups.

**Fig. 3:**
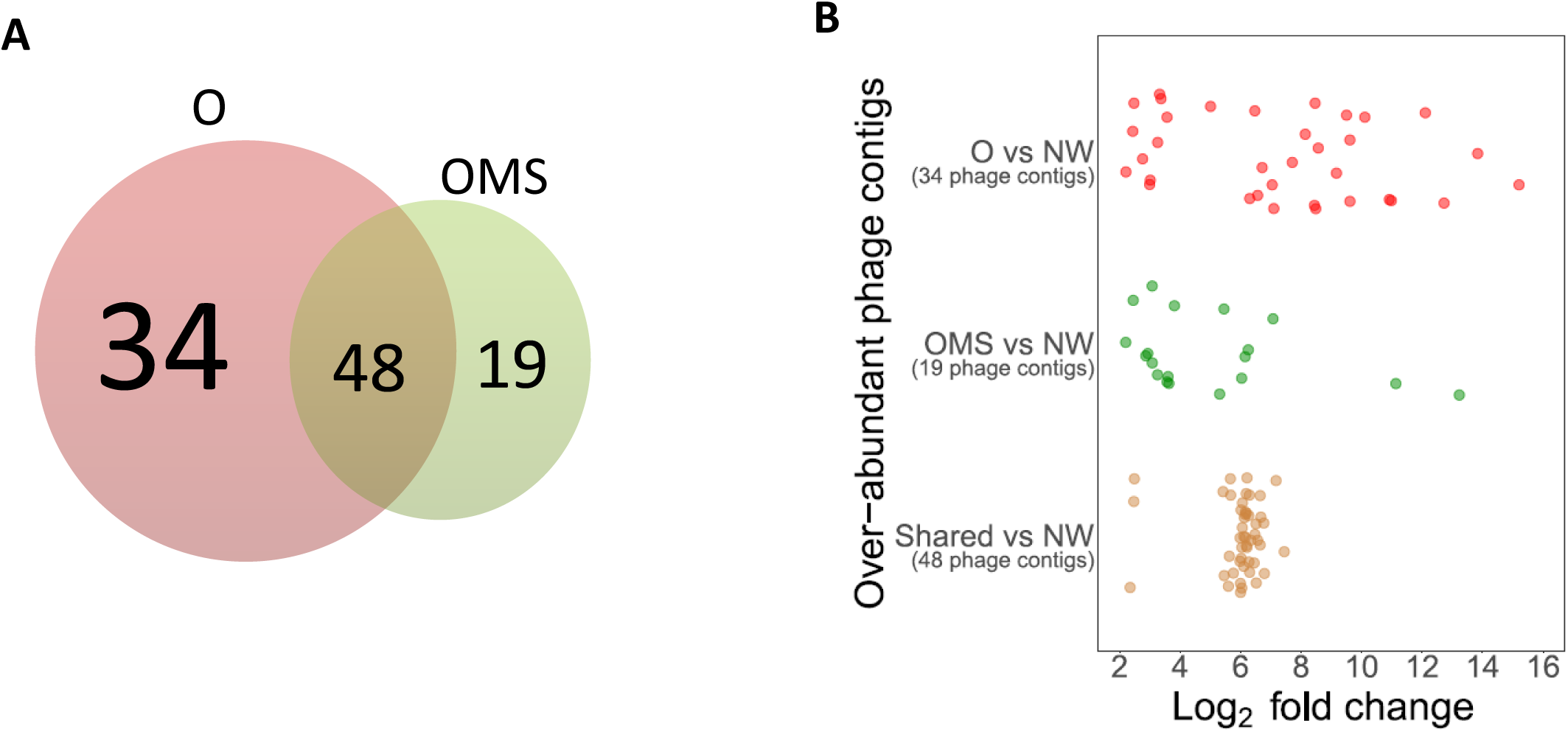
Over-abundant phage contigs in obesity. **A**. Venn diagram of the number of phage contigs that were over-abundant and present in >30% of individuals within the O group (green circle) and the OMS group (red circle), as compared with NW. The numbers of over-abundant shared phage contigs between O and OMS groups are shown in the brown area. **B. The expression levels of over-abundant phage contigs** according to their log2 fold change values in O (red points), OMS (green points), and shared (brown points), compared to the NW group. Over-abundant phage contigs were selected by a log2 fold change ≥2 and an FDR adjustment (p-value < 0.05). The abundance matrix was normalized by the RPKM method.

### Phageome richness and diversity is increased in obesity groups

To assess the relationship within-sample and between-sample phageome diversity, we examined α-diversity and β-diversity, respectively. The α-diversity metrics showed that phage richness and diversity significantly increased in the obesity groups compared to NW (Fig. 4A and B). The O group exhibited higher richness followed by OMS and NW (Fig. 4B), while the OMS group exhibited higher diversity followed by O and NW (Fig. 4A). The between-sample diversity comparison observed in a principal component analysis (PCoA) based on Bray-Curtis distances showed no defined clusters by group (Fig. 4C). Interestingly, when all the obesity samples were tagged together (O + OMS), obesity samples tend to cluster separately from NW samples (Fig. 4D).

**Fig. 4:**
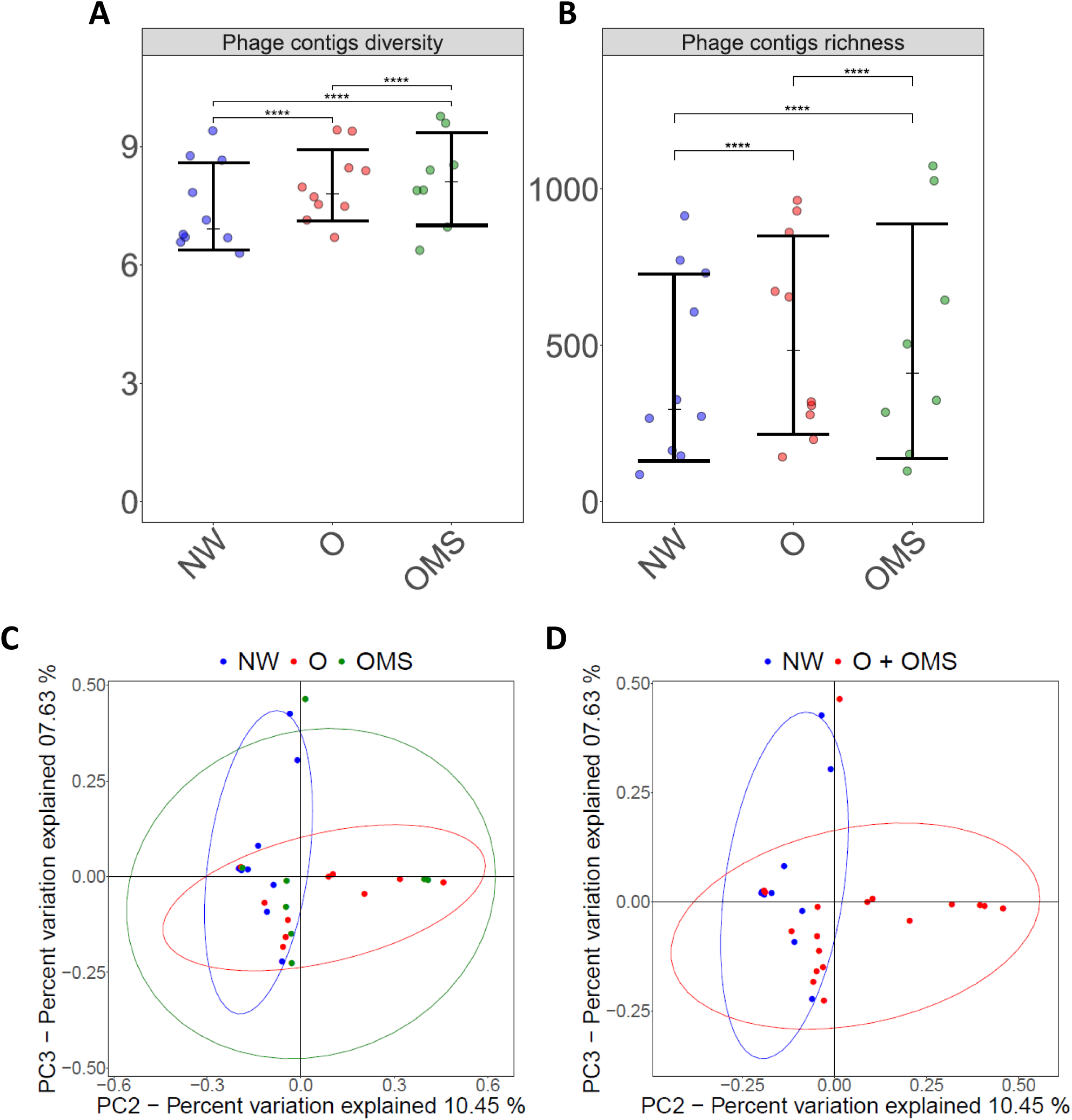
Alpha and Beta diversity of the phageome. **A**. Phage contigs diversity. The average of Shannon’s index values from 10,000 rarefactions per sample is shown as points. Error bars represent the mean +/- SD per group. **B**. Phage contigs richness. The average of the number of unique phage contigs from 10,000 rarefactions per sample is shown as points. Error bars represent the mean +/- SD per group. Statistical significance was determined by the Kruskal-Wallis test with Dunn’s correction comparing all samples to all samples. ****= p < 0.0001. Richness and diversity were calculated over the RPKM matrix. **C and D**. Principal Coordinates Analysis (PCoA) based on Bray-Curtis dissimilarity. Elipses were calculated based on the most distant samples. **(C)** Samples tagged as NW, O, and OMS: and **(D)** samples tagged by all obese (O + OMS) and NW.

### Few phage contigs dominated the phageome composition

We observed that 488 (10.58%) phage contigs accounted for 70% of the normalized reads in the NW samples, while 679 (14.73%) and 831 (18.02%) phage contigs, accounted the 70% of the normalized reads in O and OMS groups. This is in accordance with the higher diversity observed in the obesity groups and suggested a considerable change in the number of dominant phage contigs in the gut microbiome, being the highest is obesity with metabolic syndrome, followed by obesity and normal weight.

### The majority of the phageome was shared among all individuals

The recruitment matrix was analyzed to assess the composition of the phageome in all the samples independently of the health status. From the 4,611 phage contigs, 2,605 (56.50%) were present in 20-50% (termed “common phages”) and 1,088 (23.56%) were present in >50% individuals (termed “core phages”) and two (0.04%) were present in all individuals. Contrary, 847 (18.37%) phage contigs were detected in 2-19% of individuals (termed “low overlap phages”), and 71 (1.54%) in only one individual (termed “unique phages”) (Fig. 5A). Interestingly, two of the “core phages” were identified as putative “crass-like” phages. The identification of “crass-like” phages supports our phageome definition of “core phages”, due to these are the most abundant phages reported in the human gut from adult individuals^34^.

**Fig. 5.**
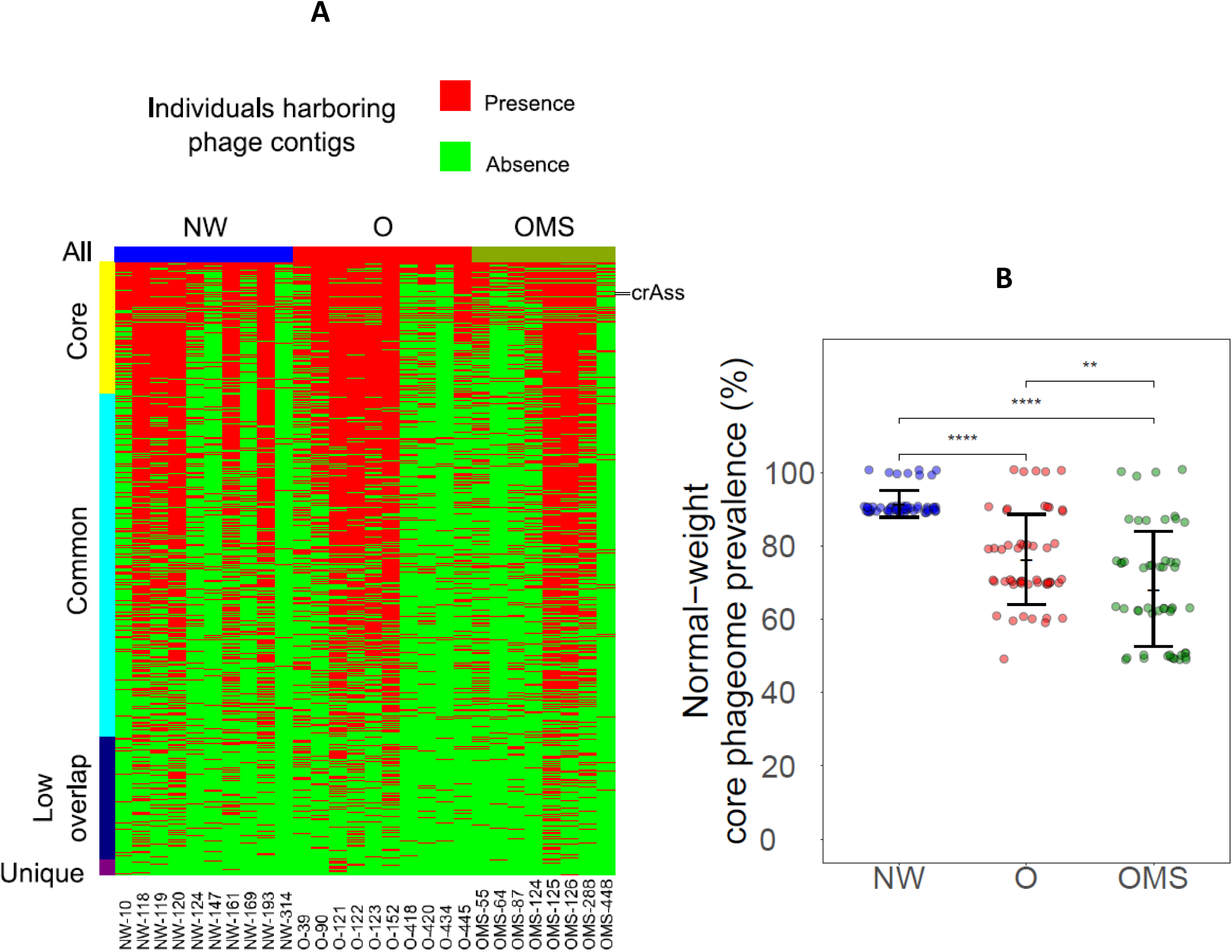
Phageome was mainly shared in the population, and the prevalence of the normal-weight core phageome was affected by obesity. Presence-absence heat map of the phage contigs distributed among all the samples. Core phages: phage contigs detected in > 50% of the individuals. Common phages: phage contigs detected in 20-50% of the individuals. Low overlap phages: phage contigs detected in 2-19% of the individuals. Unique phages: phage contigs appearing in only one individual. The core phage contigs include two phage contigs classified as crAssphage. Each line represents one phage contig. **B**. Reduction of the normal-weight core phageome (phage contigs present in >80% of NW samples) in obese children. The percentages of the 52 phage contigs’ prevalence that conform to the normal-weight core phageome are indicated by points, showing the distribution within each group. Error bars represent the mean +/- SD per group. Statistical significance was determined by the Kruskal-Wallis test with Dunn’s correction comparing all samples to all samples. ****= p < 0.0001, **= p< 0.05.

### Abundances of core phage contigs correlate with gut bacteria altered by obesity

We calculated the Spearman correlation between the 48 phage contigs present in ≥80% of all the samples and 41 bacterial taxa significantly altered among NW, O, and OMS, determined in a previous study from our group in the same set of samples, resulting that the abundance of four phage contigs (Extended Fig. 3A) correlated with bacterial taxa (Fig. 6A). Interestingly, phage contig2740, a potential Myoviridae, which was significantly more abundant in O (Extended Fig. 3A), showed a positive correlation with *Collinsella aerofaciens* (Fig. 6A), an over-abundant bacteria in the OMS group. The phage contig313, a potential member of Siphoviridae, which was significantly over-abundant in O (Extended Fig. 3A), showed a positive correlation with *Parabacteroides distasonis* (over-abundant in O Vs. OMS), and a negative correlation with an undetermined specie from genus Phascolarctobacterium (over-abundant in NW Vs. O) (Fig. 6A). The phage contigs 207 and 540, members of the Siphoviridae family, had the lowest abundance in OMS and O compared to NW (Extended Fig. 3A) and were negatively correlated with Erysipelotrichaceae, an over-abundant family in O and OMS groups (Fig. 6A). These data suggest that changes in the abundances of specific phage-bacteria interactions could be associated with obesity and may promote signatures of the phageome-bacterial interactions between obesity and obesity with metabolic syndrome.

**Fig. 6:**
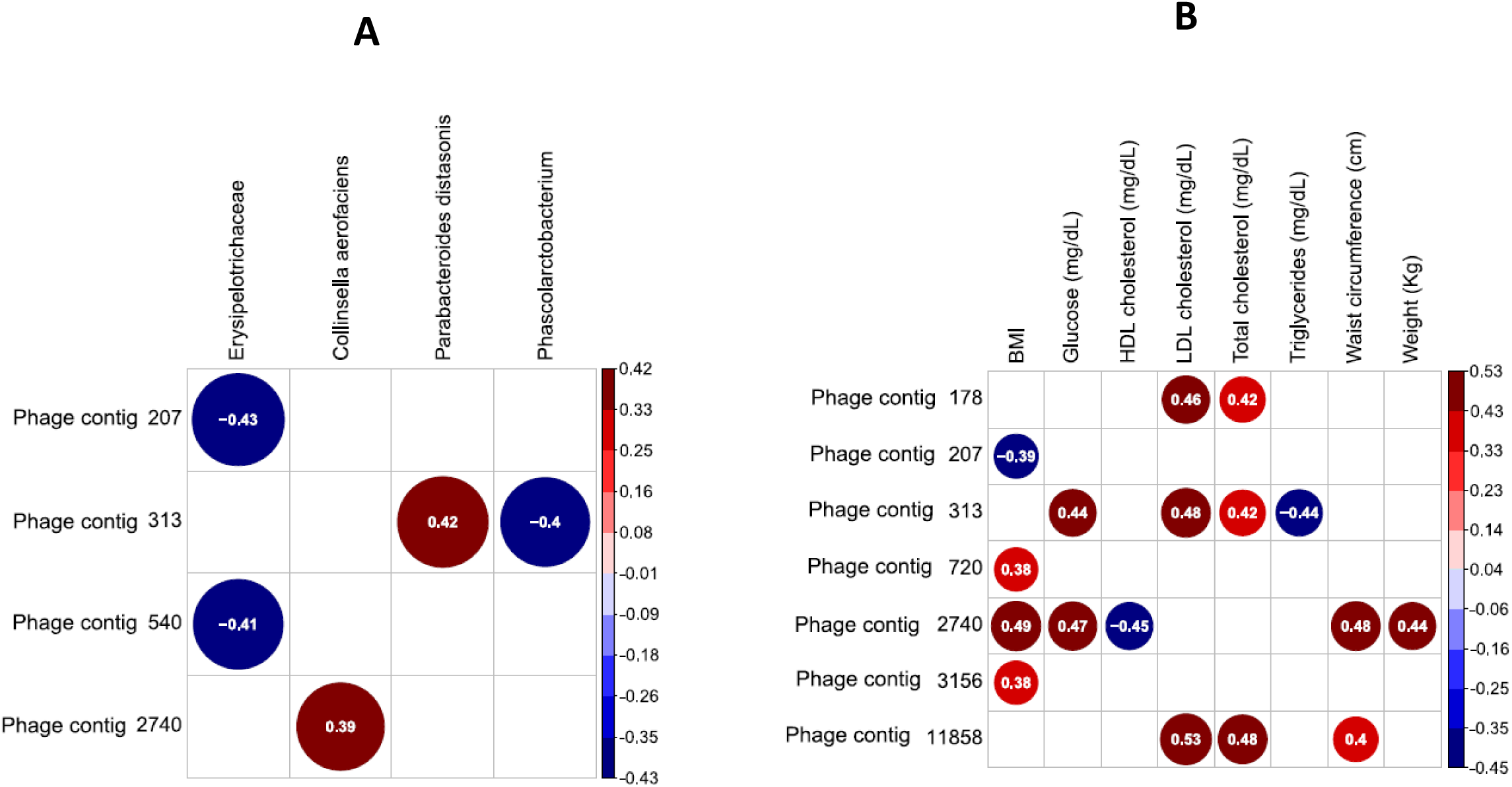
Obesity-specific changes in the enteric phageome. Spearman correlation plots of the abundances (RPKM) of the most abundant phage contigs and the relative abundance of bacterial taxa identified to be significantly associated with disease (A), and the clinical and anthropometrical parameters of obesity and metabolic syndrome (B). Positive values (red circles) indicate positive correlations, and negative values (blue circles) indicate inverse correlations. The circles’ shading indicates the magnitude of the correlation where darker shades are more correlated than lighter shades. Statistical significance was determined for all pair-wise comparisons; only significant (p-value < 0.05) are displayed. Phage contigs selection criteria for correlations: present in at least 80% of total samples.

### Abundances of shared phage contigs correlated with anthropometric and biochemical parameters altered by obesity

The Spearman correlation between the 48 phage contigs present in ≥80% of all the samples and the anthropometrical and biochemical parameters showed positive correlations between the abundance of several phage contigs (Extended Fig. 3B) and LDL levels, total cholesterol, glucose, waist circumference, BMI, and weight (Fig. 6B). In contrast, we also found a negative correlation between the abundance of several phage contigs (Extended Fig. 3B) and BMI, HDL levels, and triglycerides (Fig. 6B). These correlations suggest an association between phage contigs and typical host affected anthropometrical and clinical parameters by obesity.

### The healthy core-phageome shifts from normal weight to obesity

To analyze if the NW core phageome was related to a good health status, we compared the prevalence of core phage contigs in NW children (n = 10) to those in O (n= 10) and OMS (n = 8). To this end we selected the phage contigs shared among >80% of the NW samples. Therefore, we established a NW core phageome of 52 phage contigs, and compared its prevalence with respect to O and OMS groups. NW core phageome was found in an average of 91.54% NW individuals, and it was significantly reduced to 76.35% (p-value= <0.0001) and 68.27% (p-value= <0.0001) on average in O and OMS patients, respectively (Fig. 5B). These results show the gut core phageome of normal-weight children was significantly altered in children with obesity, being most affected in in children with obesity with metabolic syndrome.

## Discussion

Microbiome changes are widely accepted to have a significant association with human health, while metagenomic analysis of viruses, one of the most poorly understood components of the human gut microbiome, has recently revolutionized our view of gut microbiome^20,38–40^, highlighting the critical role of the interrelationships between gut phages and bacteria in health and disease.

We report a viral metagenome with one of the largest^37^ sequencing depths, ∼75M of viral reads, analyzing the phageome of school-age children with obesity and obesity with metabolic syndrome and comparing to normal-weight controls. Instead of solely using the sequencing reads, we used a *de novo* approach to construct the phageome community, composed of 4,611 phage contigs, representing complete or partial genomes. Unlike most of the human gut viromes reported to date (except Manrique et al., 2016^37^), our approach was not influenced by PCR whole-genome over-amplification that can preferentially amplify small circular ssDNA viruses, e.g. the Microviridae family^41^. In contrast, we used a tagmentation (TAG) method that allows the use of low input DNA and its free of PCR over-amplification, although it may select against ssDNA templates^41,42^. Intriguingly, contrary to the high abundance of Microviridae found in most virome studies^26,35,43,44^, this family was found at a very low abundance in our samples, probably, as a direct result of the TAG method selected for this study. This ambiguity highlights the need for quantitative analysis protocols in virome studies, as recently described^41^.

For the first time, we showed that children with obesity and obesity with metabolic syndrome had significant and specific changes in their gut phageomes. Importantly, the recruited children came from similar lifestyles, the same ethnographic region and relatively homogeneous environments, making the effects of socioeconomic, cultural, and nationality not confounding factors. We consider this feature is important to minimize bias in our results, as reported in previous reports^45–48^.

First, we observed the same number of VLPs in all the samples, independently of the health status. Contrary to previous reports on IBD^26^, ulcerative colitis^26^ and Crohn’s disease^26^. Still, we found a significant increase in diversity and richness in OMS and O, suggesting that the expansion of specific phages could be associated with the decreasing of others maintaining the same number of VLPs in the gut. Similar observations in richness and diversity have been reported in Crohn’s disease and ulcerative colitis^26^ and, a colitis murine model^49^. In contrast, a low phage richness, and diversity were detected in individuals with type I diabetes mellitus^24^. Caudovirales, was the most abundant type of phages independently of the obesity phenotype, and regardless of sequencing reads or metagenomic assembly. The predominance of this order is consistent with previous reports of the human gut viromes^26,31,37^.

We detected a low inter-individual variability of phageome. Only ∼20% of phages were low-overlap or unique, while ∼80% represented common and core phages, independently of the disease. In this regard, we also identified that the prevalence of the core phageome that was shared in >80% of the normal-weight individuals was significantly reduced in obesity, an even more in OMS. This suggests that the loss of some normal-weight phage contigs in obesity could play a role into encouraging the children’s progression to obesity. Our observation is in agreement with previous studies where a core phageome was proposed for a set of healthy humans and; also, it was reduced in patients with inflammatory bowel disease^37^.

The concept of a core virome has also received support from a recent study with adult monozygotic twins, in which 18 contigs were found to be present in all individuals^32^. In contrast to these findings, the compilation of a large-scale gut virome database called into question of the existence of a human core gut virome^19,50^. This disparity is mainly due to the well-established belief that the human gut virome is highly individual-specific^19,32,35,51^ and because the criteria used to define the presence of a viral sequence in a sample^50^ is still questionable. Despite the difficulties, we speculate, according to the previous reports^45,47,52^, that the highly homogeneous geographic and ethnic representation across our dataset were essential factors that allowed us to establish a core phageome. Additionally, we followed similar sequence thresholds recently proposed for accurate estimation of viral community composition and diversity^53^, such as (i) contig lengths ≥4 kb, (ii) coverage determined from reads mapped at ≥90% identity, and (iii) ≥80% of contig length with ≥1× coverage. However, it should be noted that our assemblies may represent fragments of the same phage genome or family. More extensive studies using stringent standard bioinformatics parameters, such as previously suggested^53^, are necessary to generate even more accurate estimates about the core phageome and its prevalence in the human population.

Our research group previously reported^18^ for the same set of samples that OMS and O groups had a significantly higher bacterial richness and diversity than NW group^18^. The same was observed in the phageome, where the OMS and O had significantly higher richness and diversity than NW. This observation is in agreement with the proposed that virome diversity correlates with intestinal bacterial diversity^32^. We also found distinct phageome and bacterial^18^ structure among OMS and O groups. Hence, considering their microbiomes, is very important that obesity and obesity with metabolic syndrome are studied as separate conditions to better understand the interplay between bacteria and phages and their role in these pathologies.

The quantitative analysis of the gut phageome allowed us to identify overabundant phage contigs in O and OMS as compared to NW, suggesting that they could be related to the physiopathology of obesity and development of metabolic syndrome. We consider that specific overabundant phage contigs in OMS could be biomarkers related to the development of metabolic syndrome in obese children, although, all potential phages must be studied in depth to generate exact conclusions and associations. It is known that alterations in the abundance of phages can also alter the abundance of specific gut bacterial species^54^. We observed that the increased abundance of phage contig 2740 positively correlated with a high abundance of *Collinsella aerofaciens*, high levels of LDL, BMI, waist circumference, glucose, and weight, and lower HDL levels.

Interestingly, *Collinsella aerofaciens* was significantly over-abundant in the OMS group compared to O and NW and showed a positive correlation with triglycerides while showing a negative correlation with HDL in the same samples ^18^. These results suggest that this phage-bacteria interaction could be strongly associated with the disease via increasing glucose, waist circumference, weight, and BMI while decreasing HDL levels. The increased abundance of phage contig-313 in O Vs. OMS positively correlated with a high abundance of *Parabacteroides distasonis*, high LDL levels, glucose, and total cholesterol. This bacteria was found enriched in O vs. OMS, and it was associated with increased LDL levels in the same cohort of our study^18^, and with weight gain and hyperglycemia in other studies ^55^. This suggests that this phage-bacteria interaction may be associated with metabolic changes in obese children. Contrary, the abundance of phage contig-313 showed a negative correlation with Phascolarctobacterium genus, which was enriched in the NW as compared to O^18^. It highlights that the increased abundance of this phage contig in obesity is directly associated to decreasing levels of Phascolarctobacterium, a possible protective bacteria against the obesity^18^.

Also, the increased abundance of phage contigs207 and 540 in NW was associated with a decreased abundance of Erysipelotrichaceae and BMI. This bacterial family was significantly increased in OMS and positively correlated with waist circumference^18^. Moreover, the increased abundance of this family has been associated with host dyslipidemia in the context of obesity, metabolic syndrome, and hypercholesterolemia^56^. It suggests that the increased abundance of these phage contigs in NW could be directly associated with decreasing levels of Erysipelotrichaceae, suggesting that both phage contigs could be used against the increased abundance of Erysipelotrichaceae in O and OMS groups^18^, opening the possibility of using these phages as a therapeutic option against the bacterial dysbiosis typically associated with obesity. Indeed, the gut phageome represents a source of individual phages with potential therapeutic applications ^57^.

Some phages only correlated with metabolic and anthropometric parameters of obesity and metabolic syndrome but not with bacteria taxa, showing that changes in bacterial abundance do not explain some disease-specific features of the phages. These correlations demonstrate that the dynamics phage-bacteria are specific between O and OMS children, and they highlight the importance of evaluating predator-prey relationships in detail, raising the possibility that changes in the abundance of specific phages and bacterial taxa may contribute to the disease. Further experiments and analysis of these phage-bacteria interactions are still necessary, e.g., in gnotobiotic mice and using shotgun sequencing, along with a better understanding of lysogenic-lytic switch in viral reproduction, would help to interpret the mechanisms of association with the disease.

Compelling evidence supports the concept that shifts in the microbial system during infancy may increase the risk for obesity later in life^58,59^. Therefore, manipulating gut microbiota using phages at an early stage might offer possibilities for the prevention and treatment of the dysbiosis associated with the obesity. Fecal microbiota transplantation (FMT) revealed that phages could be co-transferred with bacteria^60^. Even, successful treatments against *Clostridium difficile* using bacteria-free fecal filtrate provided the first evidence that phageome manipulation may be an effective therapeutic strategy to stabilize the eubiosis of the bacteria in the microbiome^61^.

The role that phages play in obesity and obesity with metabolic syndrome is critical to understanding their contribution to the microbiota dysbiosis. We believe that our study provides a better knowledge of the phage-bacteria dynamics of the gut microbiome. We propose studying specific overabundant phages in O and OMS as possible biomarkers related to the development of metabolic syndrome in obese children. Also, the development of in vivo models to test the phage-bacteria dynamics in obesity will undoubtedly be an essential area that could help to complement the understanding of microbiome dysbiosis associated with obesity. A better knowledge of the intestinal phageome composition and its interaction with the microbiota and immune host system will allow us to diminish the prevalence of childhood obesity and metabolic syndrome.

## Methods

### Viral-like particles (VLP’s) isolation

Fecal samples and the biochemical parameters information used in this study were obtained as part of a previous study of our group^18^. Viral like particles (VLPs), were isolated from ∼250mg of fecal sample suspended by vortexing in 40 ml of SM Buffer (pH 7.5) previously sterilized by temperature and filtered through 0.22 µm (Nalgene). The homogenates were centrifuged at 4,700 × g for 30 min at room temperature and the supernatant was filtered through 0.45 µm (Nalgene) and 0.22 μm PES filters (Nalgene) to remove residual host and bacteria cells. The filtrate was then concentrated to 200 ul at 4°C with an Amicon Ultra 15 filter unit, 100 KDa (Millipore). The concentrated solution of VLP’s was suspended again in of SM buffer and concentrated to 200 µl at 4°C. Then, we added 40 µl of chloroform and incubated for 10 min at room temperate to degrade any remaining bacterial and host cell membranes. Free DNA in the solution was degraded with DNase following the manufacture’s procedure (Invitrogen cat No. 18047-019). The final samples were stored at -80°C until further processing.

We used epifluorescence microscopy to quantify the isolated VLPs. Briefly, we stained 10 µl of the concentrated samples with 2µl of SYBR Green (Invitrogen) and 10 µl of paraformaldehyde previously filtered through 0.22 μm PES membranes (Millipore). Three fields per sample were observed and quantified with Olympus FV1000 Multiphoton Confocal Microscope. Additionally, 8 µl from the concentrated VLPS samples were observed in the transmission electron microscopy.

### VLPs shotgun sequencing

The DNA of VLPs was extracted following the manufacture’s protocol for the QIAampMinElute Virus Spin Kit (QIAGEN). The resulting DNAs were quantified (Qubit) for each sample and were diluted with ribonuclease free water to a concentration of 0.3 ng/µl. Each sequencing library was prepared following Illumina Nextera XT DNA Library Preparation protocol with a unique barcode combination per sample following manufacturer’s recommendations. Briefly, we mixed 5 µl of DNA at 0.3 ng/µl with the tagmentation reaction mix. Next, we added the indexed oligos and amplified the library for 12 cycles. The library was purified with AMPure XP beads to obtain DNA fragments > 500 kb. The size and quality of each library was assessed with a DNA bioanalyzer 2100 (Agilent). All libraries were pooled together and sequenced using the Illumina NextSeq500 platform in the 2×150 pair-end mode at the Sequencing Unit Facility of the National Institute for Genomics Medicine, México.

### Analysis of the sequenced reads

Total reads were dereplicated. Adapters and low-quality bases (PHRED Q30) were trimmed using Trim_Galore (https://github.com/FelixKrueger/TrimGalore) and fastq toolkits tools, removing (http://hannonlab.cshl.edu/fastx_toolkit/index.html) and the first 20 nucleotides. Human and bacterial reads were removed by read mapping using BWA^62^ (against the Homo sapiens v38 reference genome) and Kraken^63^ against bacteria NR database, with default parameters. Quality filtered reads were clustered at 95% identity level using CD-HIT (http://weizhongli-lab.org/cd-hit/) to remove redundancy and to generate a unique sequence dataset. The viral richness between groups (NW, O, OMS) was determined collecting 1,000 random sub-samples of 149,000 sequencing reads according to the smallest sample (NW_169: 149775) using CD-HIT.

### Viral reads classification

Unique sequences were taxonomically classified into orders and families, according to the International Committee on Taxonomy of Viruses (ICTV), using BLastX (E-value <10-6) against the NR refseq viral database and considering the lowest-common ancestor algorithm in MEtaGenomeANalyzer (MEGAN5)^64^. Absolute read counts for selected viral taxa were normalized obtaining the relative abundance for each sample using R.

### De Novo contig assembly and taxonomic profiles of viral genomes

Total viral reads from all samples, were pooled for de Novo assembly using IDBA-UD assembler ^65^ with a k-mer length of 20-125 with scaffolding rounds. Each sample was re-mapped separately with Bowtie2 against the viral assembly using the end-to-end mode. Viral scaffolds covered ≥80% of their length in at least one sample were used as evidence of phage presence. Next, we kept the scaffolds ≥4 Kb for downstream analysis, to decrease the probability of selecting gut eukaryotic viruses (mainly Anelloviridae family) (maximum reported genome size approximately 3-4Kb)^31^ and considering the shorter phage genomes reported have around ≥ 4Kb^33^. To eliminate the redundant contigs we used CD-HIT using 95% clustering identity. The taxonomy annotation of each contig was obtained using dc_megablast against the NT NCBI viral genomes database with E-value of 1×10^−6^ and maximum number of target sequences to report set to 25 hits. Using the megablast results, the taxonomy of each contig was assigned by the lowest-common-ancestor algorithm in MEtaGenomeANalyzer (MEGAN5). After that, all the contigs without taxonomic classification were used for search against the NR NCBI viral protein database using BLASTp with maximum e-value cutoff 1×10^−6^and maximum number of target sequences to report set to 50. Using the BLASTp results, the taxonomy of each gene was assigned by the lowest-common-ancestor algorithm in MEtaGenomeANalyzer (MEGAN5).

### Contigs abundance and statistical analysis

The reads recruitment to the assembly was used to construct an abundance matrix, applying the filter of coverage and length as previously recommended^53^. The coverage was defined from reads mapping (Bowtie2) at ≥90% identity and ≥80% of contig length with ≥1X coverage. Mapping outputs were converted into an abundance matrix normalized by Reads Per Kilobase per Million sequenced reads per sample (RPKM)^31^. This matrix was used to determine statistical differential contig-abundance using edgeR with log_2_ fold change ≥2 and FDR adjustment (p-value≤ 0.05). Richness and diversity were calculated over the RPKM matrix using QIIME 1.9^66^at 10,000 rarefactions.

### Phage correlations with bacteria and biochemical parameters

All correlations were calculated using the Spearman coefficient with R. For the abundance of phages, we considered the RPKM matrix and for the microbiota, we used the relative frequency of the significantly taxa previously reported between NW, O, and OMS^18^.

### Data accessibility

The assembly of phage contigs was deposited in the GenBank under the BioProject SUB7771982.

## Declarations

### Ethics approval and consent to participate

The Ethic Committee of the National Institute of Genomic Medicine (INMEGEN) in Mexico City approved the study. The parents or guardians of donors signed the informed consent form for participation, and the donors assented to participate.

### Consent for publication

Not applicable.

### Availability of data and materials

All data generated during this study are included in this published article. The sequencing data have been deposited in the SRA repository of NCBI and can be accessed via the BioProjectSUB4772779. Requests for material should be made to the corresponding authors.

### Authors’ contributions

Conceptualization: SB, GLL, FCG and AOL; Data curation: SB, GLL, FCG, LGB, FS, EEM, JPOR, BELC, SCQ and AOL; Formal analysis: SB, GLL, FCG, LGB, FS, EEM, JPOR, BELC, SCQ and AOL; Funding acquisition: SCQ and AOL; Investigation and Methodology, SB, GLL, FCG, LGB, FS, EEM, JPOR, and AOL; Project administration: SB, GLL, FCG and AOL; Writing original draft: SB, GLL, FCG and AOL; Writing review & editing: SB, GLL, FCG, LGB, FS, EEM, JPOR, BELC, SCQ and AOL. All authors read and approved the final manuscript.

## Acknowledgements

We thank Juan Manuel Hurtado Ramírez for the informatics technical support and to Abigail Hernández-Reyna for his helpful support on molecular biology protocols. We thank the National Laboratory Advanced Microscopy in the Biotechnology Institute (IBt) at UNAM for help with epifluorescence microscopy technical support and Dr. Guadalupe Trinidad Zavala Padilla at UNAM for her help with TEM microscopy. S.B. thanks to the Master’s and Doctoral Program in Medical, Dental and Health Sciences, in the field of knowledge: Experimental Clinical Research in Health, disciplinary field: Clinical Biochemistry, at UNAM. Doctoral fellowship: 405490260, CVU: 481385. F.C.G., and L.G.B., acknowledges the support of CONACyT as a Postgraduates fellows. G.L.L. was supported by CONACyT postdoctoral fellowship CB-2013/223279. We also thanks to Alfredo Mendoza-Vargas of the Unidad de Secuenciación Masiva from INMEGEN for sequencing technical support. There is not a conflict of interest for authors.

## Figure legends

**Extended Fig. 1. VLPs sequences detected in NW, O, and OMS samples**.

Plots of the total number of sequences (A), quality-filtered sequences (B), and human and bacteria quality-filtered sequences (C). Error bars indicate the median and interquartile range. The number of sequences per sample is shown as points. There were no statistically significant differences among groups.

**Extended Fig. 2**.

Virus taxonomic assignment of **VLPs derived reads. A**. The relative abundance of unique sequences matched against a viral protein sequence (hits), or not matched (no hits), is shown for each group. **B. The r**elative abundance of unique sequences assigned to Caudovirales and their taxonomic family members in NW, O, and OMS groups.

The number of unique sequences per sample is shown as points. Error bars indicate the median and interquartile range.

**Extended Fig. 3:** Box plots of the phage contig abundances in NW, O, and OMS groups that significantly correlated with **A**, bacterial taxa altered by obesity and obesity with metabolic syndrome and **B**, the clinical and anthropometrical parameters of obesity and metabolic syndrome. The phage contigs 2740, 313, and 207 showed in **“A”** also correlated with clinical and anthropometrical parameters. Points represent the phage contig abundances (RPKM) per sample. Error bars indicate the median and interquartile range per group.

