## Supplementary figures and images for "Gut Phageome Analysis Reveals Disease-Specific Hallmarks in Childhood Obesity"

### Extended Figure 1

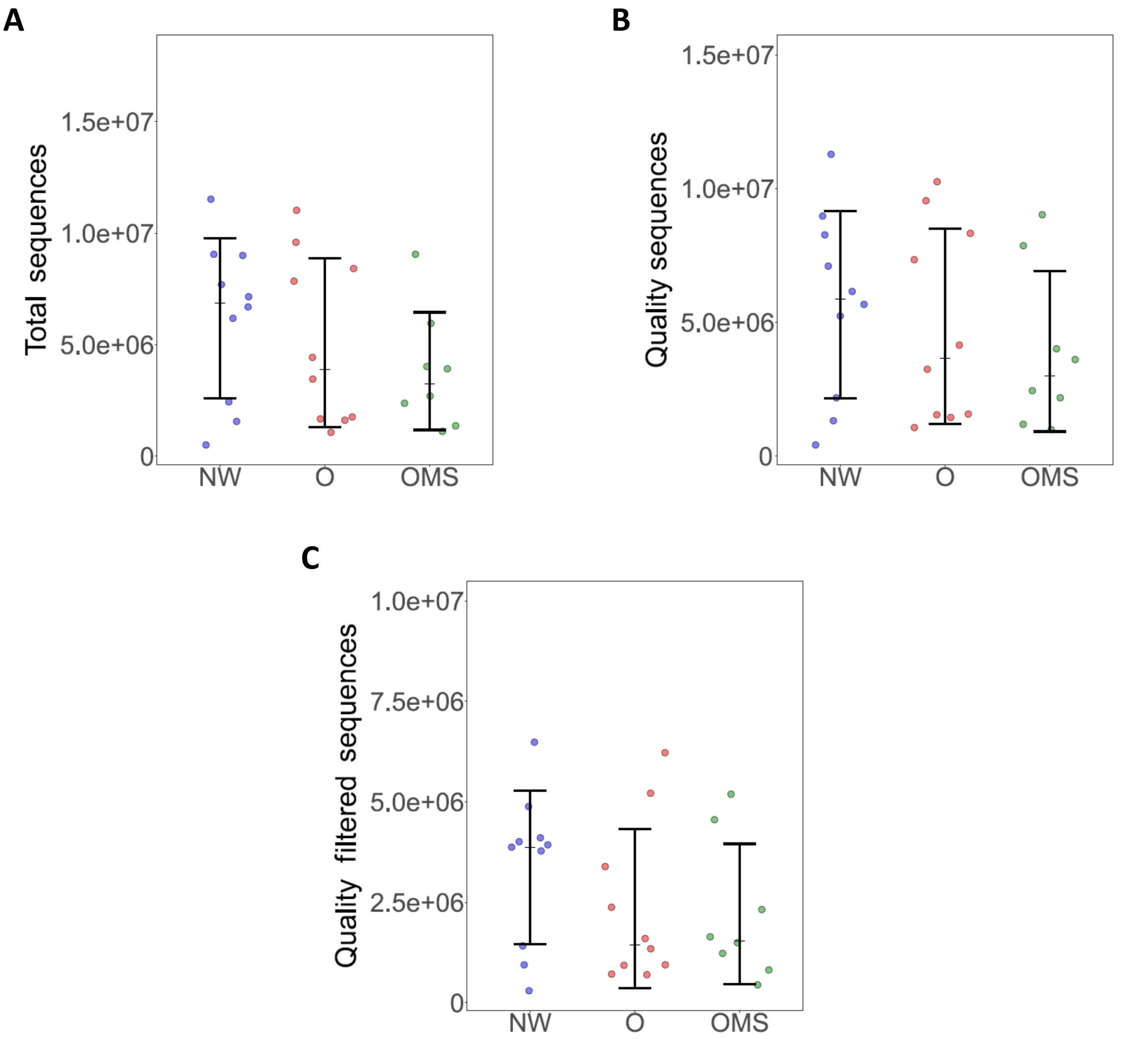


**Extended Fig. 1**

### Extended Figure 2

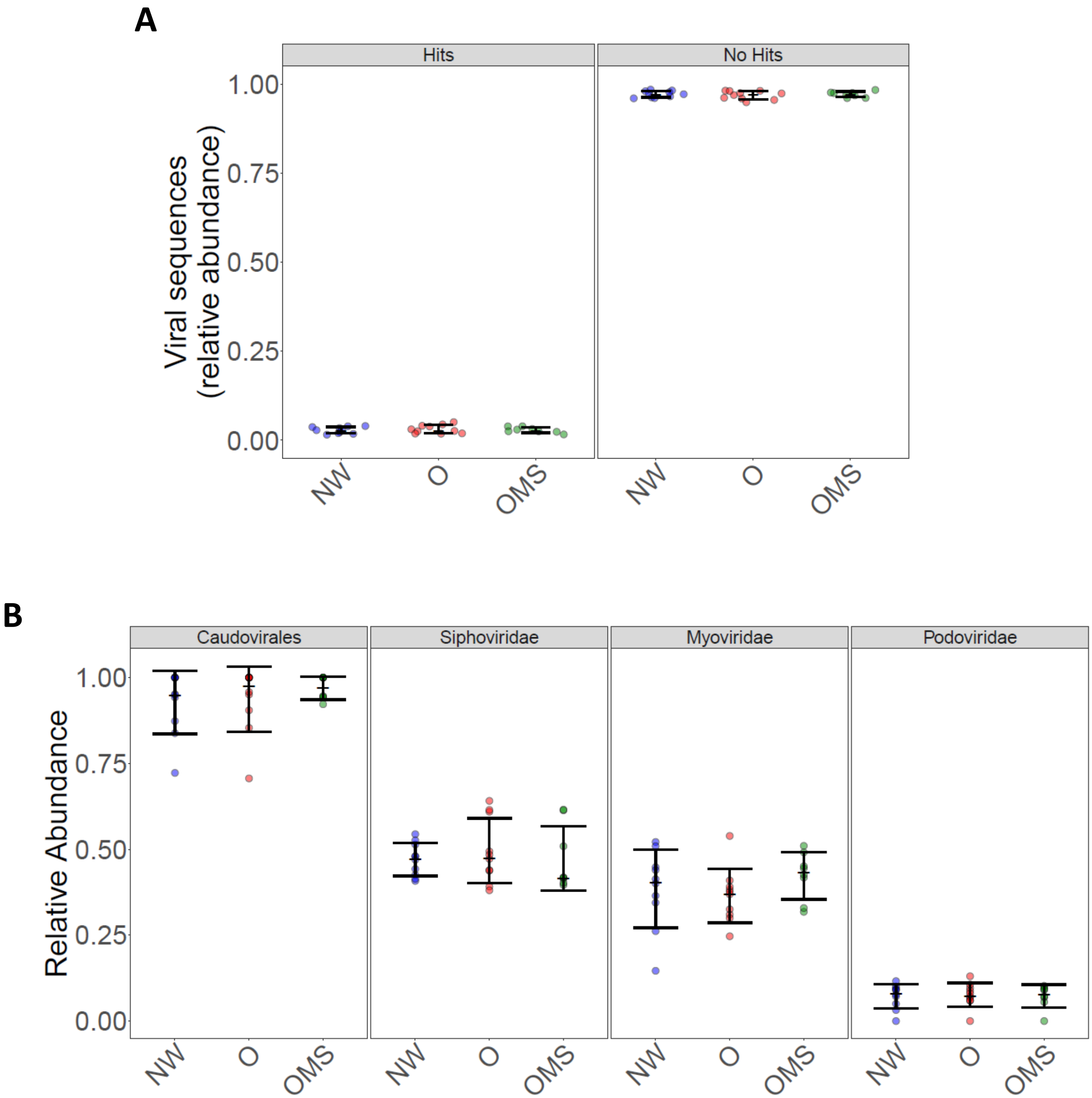


**Extended Fig. 2**

### Extended Figure 3

A

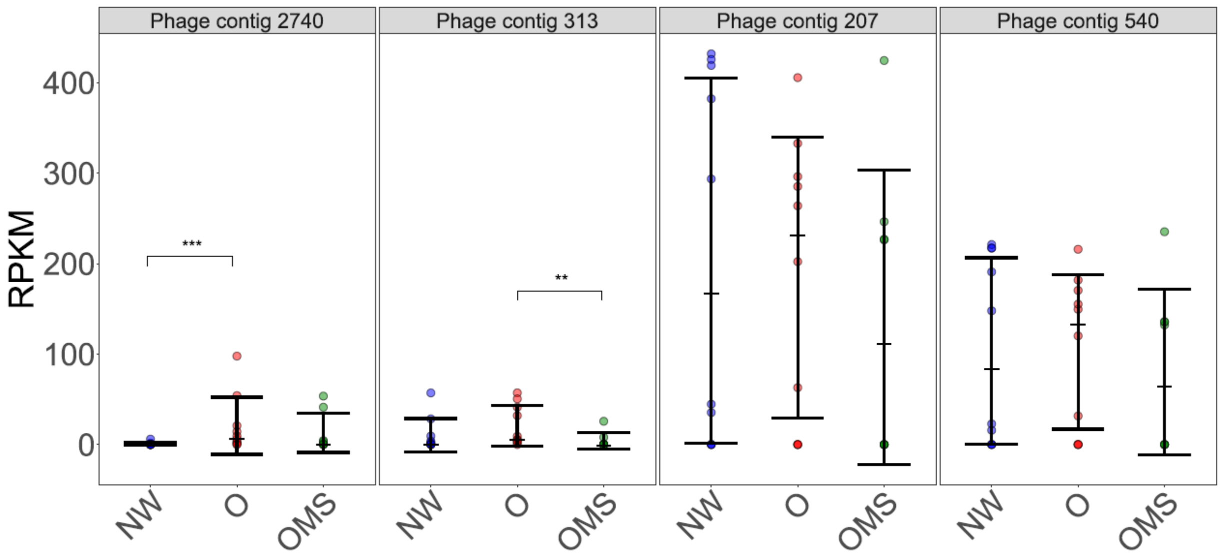


B

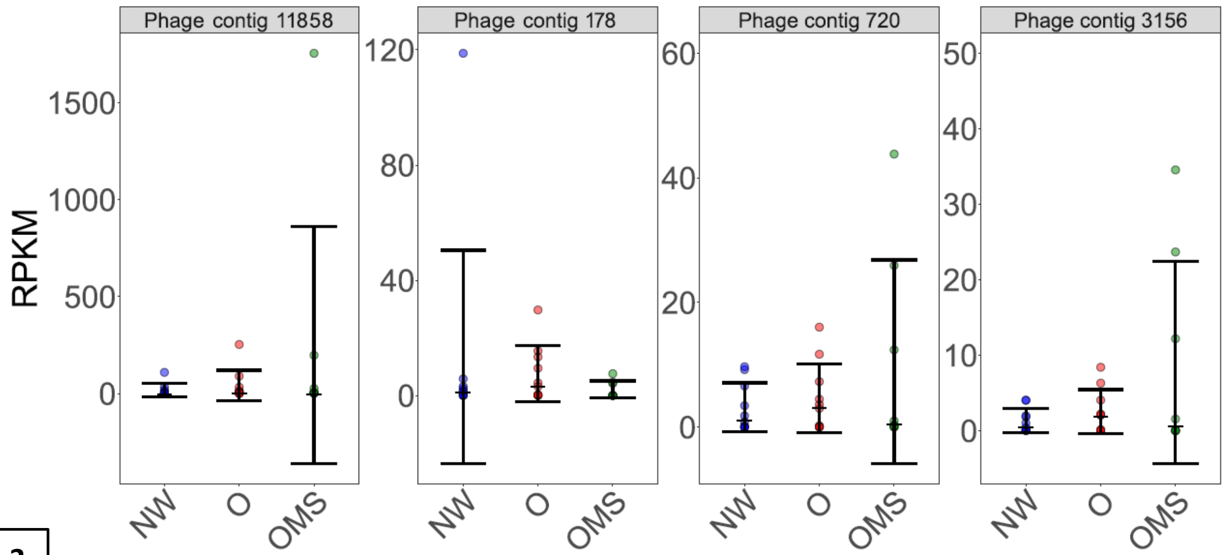


Extended Fig. 3
