## Extended Table 1 for "Gut Phageome Analysis Reveals Disease-Specific Hallmarks in Childhood Obesity"

| Samples ID | Gender | Age | Weight (Kg) | Height (cm) | BMI (percentile) | WC (cm, percentile) | BP (mmHg, percentile) | Glucose (mg/dL) | TG (mg/dL) | HDL-cholesterol (mg/dL) |
|---|---|---|---|---|---|---|---|---|---|---|
| **NW_118** | Female | 9 | 34 | 142 | 56th | 58.5, 25th | 95/70, 23th/79th | 81 | 79 | 50 |
| **NW_119** | Male | 7 | 26 | 128.6 | 58th | 56.3, 25th | 98/65, 40th/69th | 92 | 35 | 61 |
| **NW_120** | Female | 8 | 29 | 134 | 56th | 59.4, 25th | 91/66, 19th/72th | 91 | 44 | 71 |
| **NW_010** | Male | 7 | 22 | 121.3 | 31th | 61.4, 25th | 89/67, 26th/80th | 81 | 95 | 50 |
| **NW_124** | Male | 10 | 32 | 141.2 | 30th | 60.9, 25th | 100/75, 38th/88th | 86 | 57 | 54 |
| **NW_147** | Female | 9 | 30 | 132.3 | 65th | 63.1, 25th | 89/60, 14th/51th | 79 | 66 | 61 |
| **NW_161** | Male | 9 | 28 | 134 | 36th | 60.4, 25th | 97/64, 38th/64th | 94 | 42 | 78 |
| **NW_169** | Male | 8 | 27 | 134 | 27th | 58.5, 25th | 99/71, 45th/82th | 92 | 71 | 50 |
| **NW_193** | Male | 9 | 22 | 122.2 | 18th | 55.1, 25th | 92/62, 38th/64th | 85 | 42 | 75 |
| **NW_314** | Female | 7 | 22 | 124.5 | 15th | 49.6, 25th | 86/61, 14th/61th | 81 | 35 | 75 |
| **OB_121** | Male | 10 | 43 | 137.6 | 95th | 80.4, >75th | 100/68, 44th/74th | 90 | 55 | 52 |
| **OB_122** | Male | 9 | 44 | 138.5 | 96th | 87, >90th | 99/62, 37th/52th | 90 | 56 | 59 |
| **OB_123** | Female | 9 | 62 | 139.9 | 99th | 100.4, >90th | 100/72, 42th/85th | 90 | 86 | 51 |
| **OB_152** | Male | 9 | 63 | 153.5 | 98th | 96.5, >90th | 113/68, 71th/64th | 90 | 73 | 39 |
| **OB_039** | Male | 8 | 40 | 137 | 95th | 76.7, >90th | 86/54, 7th/27th | 97 | 90 | 30 |
| **OB_418** | Male | 10 | 51 | 148.6 | 96th | 80, > 75 th | 117/57, 84th/32th | 91 | 64 | 41 |
| **OB_420** | Female | 7 | 28 | 119.9 | 95th | 71, >90th | 82/61, 10th/63th | 91 | 93 | 52 |
| **OB_434** | Female | 8 | 38 | 135.7 | 95th | 70.7, > 75 th | 105/59, 64th/46th | 88 | 151 | 54 |
| **OB_445** | Male | 8 | 43 | 137.3 | 98th | 77.3, >90th | 120/79, 95th/94th | 93 | 60 | 52 |
| **OB_090** | Male | 9 | 45 | 143.2 | 95th | 84.4, >90 th | 91/64, 11th/57th | 96 | 72 | 40 |
| **OMS_124** | Male | 9 | 59 | 146.1 | 98th | 91.9, >90th | 124/80, 96th/93th | 81 | 135 | 47 |
| **OMS_125** | Male | 7 | 52 | 136.4 | 99th | 90.2, >90th | 98/62, 32th/54th | 89 | 152 | 33 |
| **OMS_126** | Female | 9 | 44 | 131.2 | 98th | 89.4, >90th | 88/60, 14th/53th | 79 | 126 | 42 |
| **OMS_288** | Male | 7 | 48 | 136.5 | 99th | 88.2, >90th | 104/72, 55th/83 | 91 | 128 | 39 |
| **OMS_446** | Male | 9 | 45 | 133.2 | 98th | 83.3, >90th | 119/78, 96th/94th | 90 | 306 | 33 |
| **OMS_055** | Male | 10 | 55 | 142.5 | 98th | 95.6, >90th | 88/60, 7th/46th | 91 | 276 | 24 |
| **OMS_064** | Female | 10 | 42 | 136 | 95th | 77.6, >75th | 99/68, 41th/76th | 84 | 331 | 31 |
| **OMS_087** | Male | 10 | 68 | 149.2 | 99th | 103.7, >90th | 100/61, 28th/45th | 91 | 121 | 28 |

**Table S1**
