## Extended Table 3 for "Gut Phageome Analysis Reveals Disease-Specific Hallmarks in Childhood Obesity"

| Dataset | N° of reads | Length (bp) | N° of total bp | n | bp per individual |
|---|---|---|---|---|---|
| Reyes | 1,318,248 | 251 | 331,454,378 | 12 | 27,621,198 |
| Minot | 907,909 | 544 | 493,947,707 | 6 | 82,324,618 |
| Norman | 153,591,468 | 250 | 38,397,867,000 | 62 | 619,320,435 |
| Manrique 1.1 | 5,813,020 | 300 | 1,743,906,000 | 1 | 4,031,007,600 |
| Manrique 1.2 | 7,623,672 | 300 | 2,287,101,600 | 1 | |
| Manrique 2.1 | 6,631,762 | 300 | 1,989,528,600 | 1 | 4,126,410,600 |
| Manrique 2.2 | 7,122,940 | 300 | 2,136,882,000 | 1 | |
| **Bikel** | **74,859,356** | **150** | **11,228,903,400** | **28** | **401,032,264** |

**Table S3**
